# The 4D Nucleome Project

**DOI:** 10.1101/103499

**Authors:** Job Dekker, Andrew S. Belmont, Mitchell Guttman, Victor O. Leshyk, John T. Lis, Stavros Lomvardas, Leonid A. Mirny, Clodagh C. O’Shea, Peter J. Park, Bing Ren, Joan C. Ritland Politz, Jay Shendure, Sheng Zhong, The 4D Nucleome Network

**Affiliations:** Program in Systems Biology, Department of Biochemistry and Molecular Pharmacology, University of Massachusetts Medical School, Howard Hughes Medical Institute, Worcester, MA 01605; Department of Cell and Developmental Biology, University of Illinois, Urbana-Champaign, IL 61801; Division of Biology and Biological Engineering, California Institute of Technology,Pasadena, CA 91125; Department of Bioengineering, University of California San Diego, La Jolla, CA 92093; Department of Molecular Biology and Genetics, Cornell University, Ithaca, NY 14853; Department of Biochemistry and Molecular Biophysics, Mortimer B. Zuckerman Mind Brain and Behavior Institute, Columbia University, New York, NY 10027; Institute for Medical Engineering and Science, and Department of Physics, Massachusetts Institute of Technology, Cambridge, MA 02139; Molecular and Cell Biology Laboratory, Salk Institute for Biological Studies, La Jolla, CA 92037; Department of Biomedical Informatics, Harvard Medical School, Boston, MA 02115; Ludwig Institute for Cancer Research, Department of Cellular and Molecular Medicine, Institute of Genomic Medicine, Moores Cancer Center, University of California San Diego, La Jolla CA 92093; Basic Sciences, Fred Hutchinson Cancer Research Center, 1100 Fairview Ave. N.Seattle, WA 98109; Department of Genome Sciences, University of Washington, Seattle WA Howard Hughes Medical Institute, Seattle WA 98109; A list of participating scientists is included in supplemental table 1.

## Summary

The spatial organization of the genome and its dynamics contribute to gene expression and cellular function in normal development as well as in disease. Although we are increasingly well equipped to determine a genome’s sequence and linear chromatin composition, studying the three-dimensional organization of the genome with high spatial and temporal resolution remains challenging. The 4D Nucleome Network aims to develop and apply approaches to map the structure and dynamics of the human and mouse genomes in space and time with the long term goal of gaining deeper mechanistic understanding of how the nucleus is organized. The project will develop and benchmark experimental and computational approaches for measuring genome conformation and nuclear organization, and investigate how these contribute to gene regulation and other genome functions. Further efforts will be directed at applying validated experimental approaches combined with biophysical modeling to generate integrated maps and quantitative models of spatial genome organization in different biological states, both in cell populations and in single cells.

## Background

The human genome contains tens of thousands of genes, and a much larger number of regulatory elements, that collectively encode the program by which we develop and function as complex organisms. Large-scale studies within the last decade have extensively cataloged these components of our genome, as well as the cell types and tissues in which they are active. For instance, the ENCODE, Roadmap Epigenome, the International Human Epigenome Project and FANTOM projects ^1–4^ annotated tens of thousands of genes and millions of candidate regulatory elements such as enhancers, insulators and promoters in the human and mouse genomes. However, our understanding of where and how these elements contribute to gene regulation, including the mechanisms by which they exert regulatory effects on specific target genes across distances of kilobases to in some cases megabases, is decidedly incomplete.

It is increasingly appreciated that the spatial folding of chromosomes and their organization in the nucleus have profound regulatory impact on gene expression. For example, spatial proximity is necessary for enhancers to modulate transcription of their target genes (e.g. ^5–7^), and clustering of chromatin near the nuclear lamina is correlated with gene silencing and the timing of DNA replication ^8,9^. Meanwhile, genome-wide association studies (GWAS) have identified large numbers of disease-associated loci, and the vast majority of them are located in distal, potentially regulatory, noncoding regions (e.g. ^10^). These GWAS results not only emphasize the importance of distal regulatory elements for genome functions, but also suggest an exciting opportunity to uncover fundamental mechanisms of disease through the mapping of long-range chromatin interactions and 3D genome organization. Therefore, to fully determine how the genome operates inside living cells, we need to understand not only the linear encoding of information along chromosomes, but also its 3-dimensional organization and its dynamics across time, *i.e.* the “4D nucleome”. Concomitantly, we must pursue deeper knowledge of the biophysical and molecular factors that determine genome organization, and how this organization contributes to gene regulation and other nuclear activities.

Genome conformation is intimately connected to the broader organization of the nucleus. Over a century of microscopy studies have revealed that the nucleus is not a homogeneous organelle, but rather consists of distinct nuclear structures and non-chromatin bodies as well as defined chromosomal regions, such as centromeres, telomeres and insulator bodies, recognized to cluster with each other and other genomic regions to define distinct nuclear compartments ^11–13^. Examples of nuclear structures include the nuclear periphery, nuclear pores and the heterochromatic compartment ^14^, while examples of nuclear bodies include nucleoli, nuclear speckles, paraspeckles, and Cajal and PML bodies. Chromosome conformation capture approaches ^15^ have yielded additional insights by characterizing chromatin folding genome-wide at kilobase-resolution ^6,16–18^. These studies show that the genome is compartmentalized in active and inactive spatial compartments at the scale of the nucleus, and within each compartment folding of chromatin fibers brings together sets of loci and regulatory elements that are otherwise separated by large genomic distances. It is possible that such interactions are at least in part related to and controlled by the formation of chromatin domains ^17,19,20^. Further evidence suggests that CTCF, the cohesin complex, and many other DNA binding proteins as well as noncoding RNAs play roles in organizing chromatin domains and long-range interactions between DNA loci ^17,21–26^. Altogether, these studies point to the genome being intricately organized within the nucleus, with this organization playing a critical role in gene regulation and activity.

The past decade witnessed significant technological innovation in the area of chromosome and nuclear structure analysis. Genomic approaches for mapping chromatin interactions, such as 3C, 4C, 5C, Hi-C, and ChIA-PET, are yielding genome-wide chromatin interaction maps at unprecedented resolution ^18^. Live cell and super-resolution microscopic approaches, combined with application of new ways (e.g. CRISPR/Cas9 –based systems) to visualize chromosome loci and sub-nuclear structures are beginning to provide detailed views of the organization and dynamics of chromatin inside single (living) cells ^27–33^. There has also been tremendous progress in modeling and analyzing chromosome structural data, producing structural models for chromosome folding which promise to provide biophysical interpretation of these often complicated datatypes ^34,35^. However, despite this progress, a comprehensive understanding of the 4D nucleome is still lacking. This is partly due to the fact that different experimental cell systems and approaches are used, which together with the absence of widely shared benchmarks for assay performance have led to observations that cannot be directly compared. Additionally, we currently have limited ability to integrate different datatypes (e.g. chromatin interaction data and imaging-based distance measurements) and lack approaches that can measure and account for cell-to-cell variability in chromosome and nuclear organization. Finally, we continue to lack mechanistic insights into the relationships between chromosome conformation and nuclear processes including transcription, DNA replication, and chromosome segregation. We believe these major gaps can be addressed by a highly synergistic, multidisciplinary and integrated approach in which groups with different expertise and knowledge, ranging from imaging and genomics to computer science and physics, work closely together to study common cell systems using complementary methods.

## Overview of the goals and strategy of the project

The 4D Nucleome (4DN) Network aims to develop a set of validated approaches to map the structures and dynamics of the genome and to relate these features to its biological activities. The Network aims to generate integrated maps and comprehensive, quantitative models of nuclear organization in diverse cell types and conditions, including in single cells. Overall, we anticipate that these efforts will lead to new mechanistic insights into how the genome is organized, maintained, expressed, and replicated, in both normal and disease states.

Specifically, the 4DN Network will 1) develop, benchmark, validate, and standardize a wide array of technologies to probe the 4D nucleome; 2) integrate, analyze, and model datasets obtained with these technologies to obtain a comprehensive view of the 4DN; and 3) investigate the functional role of various structural features of chromatin organization in transcription, DNA replication and other nuclear processes. These three main components are illustrated in Figure 1.

**Figure 1.**
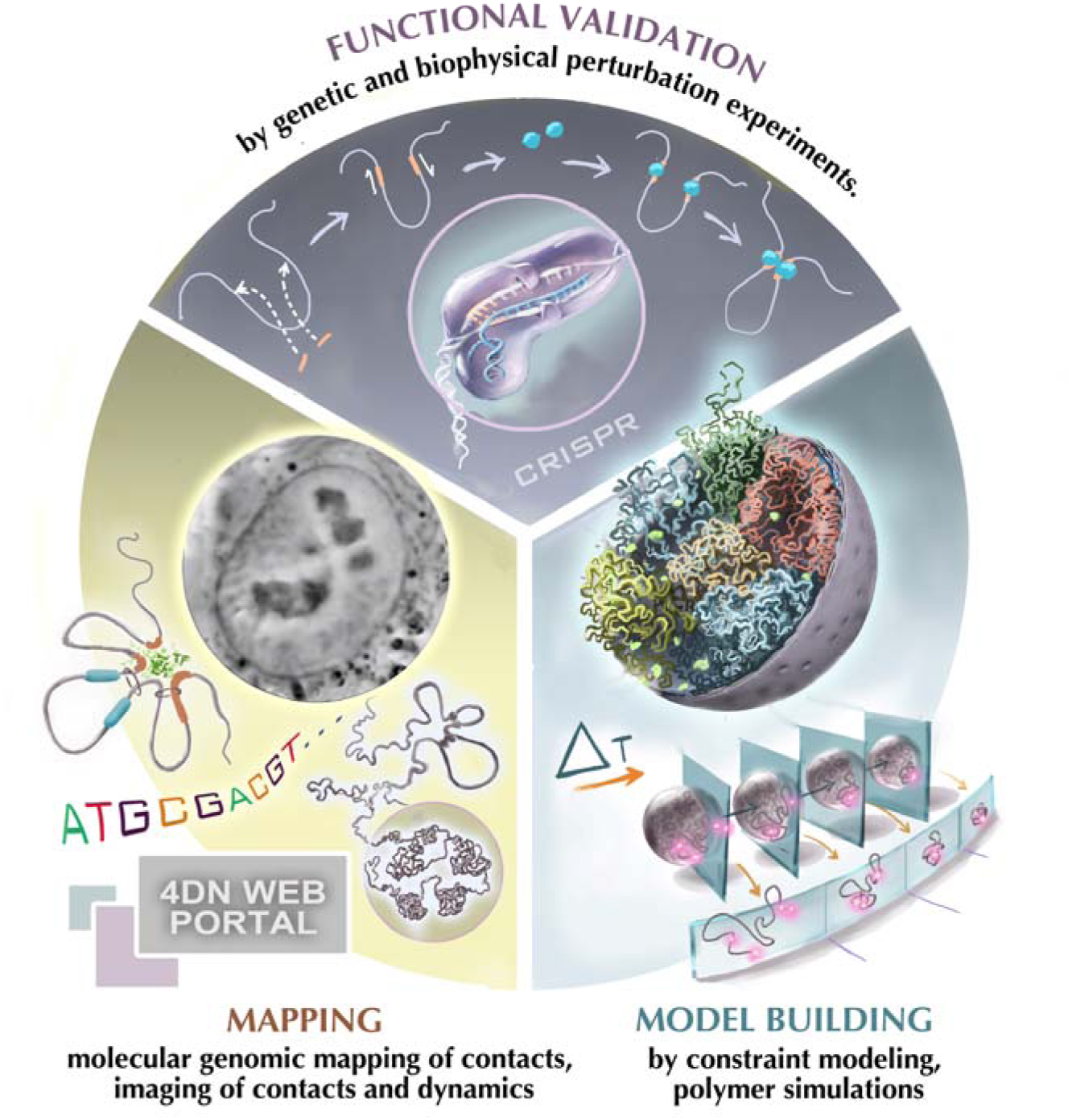
The 4D Nucleome project. The project encompasses three components: First, experimental mapping approaches are employed to measure a range of aspects of the spatial organization of the genome including chromatin loops, domains, nuclear bodies etc. Second, computational and modeling approaches are used to interpret experimental observations and build (dynamic) spatial models of the nucleus. Third, perturbation experiments, e.g. using CRISPR/cas9-mediated genome engineering, are used for functional validation. In these studies chromatin structures are altered, e.g. removing chromatin loops or inducible creating novel loops at defined positions. These perturbation studies can be complemented with functional studies, e.g. analysis of gene expression to assess the functional implications of chromatin folding. (Picture of cell nucleus was provided by Hanhui Ma and Thoru Pederson).

To achieve these objectives, we have identified the following key steps. First, a set of common cell lines will be studied across the Network to enable direct cross-validation of data obtained with different methods (Table 1). Important criteria for selecting a set of common cell lines include a stable, haplotype-phased and normal karyotype, ease of growth, ease of genome editing, and suitability for (live-cell) imaging. Further, given that cell populations are characterized by significant cell-to-cell variation in their biological state (e.g. cell cycle stage), it will be important to employ clonal cell populations that can be synchronized, activated, induced or differentiated in a controlled manner.

**Table 1.**
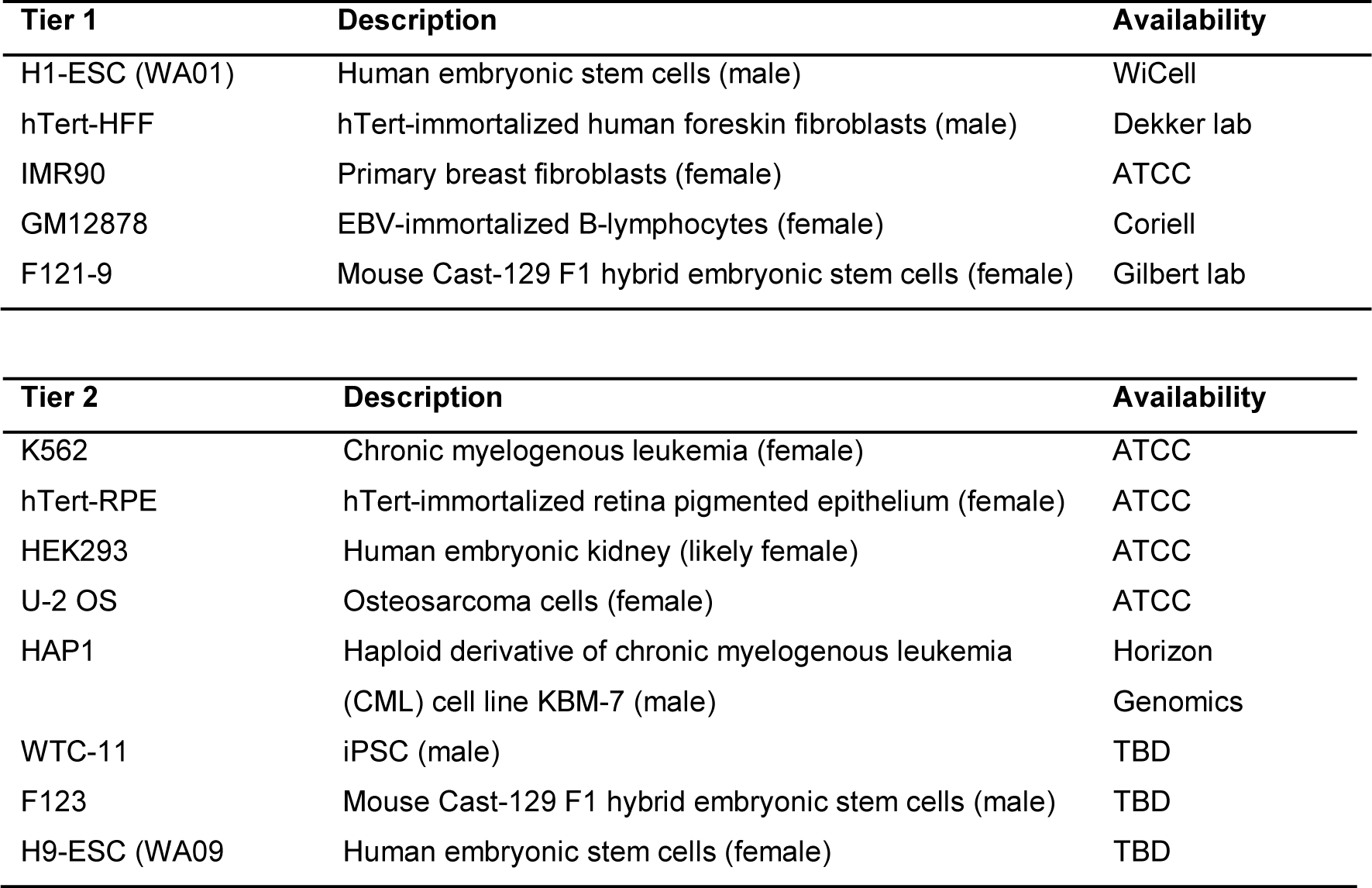
Common cell lines used by the 4D Nucleome network. Tier 1 cells are used for all studies, while Tier 2 cell lines are used for specific projects.

Second, standards for data formats and data quality will be established so that data can be shared and compared among members of the consortium and the scientific community. This includes defining metrics for reproducibility across technical replicates (for technology development) and across biological replicates, and an objective assessment of the sensitivity, specificity, resolution and precision with which aspects of the 4D nucleome can be measured.

Third, computational and analytical tools will be developed to analyze individual datasets and to integrate, compare and cross-validate data obtained with different technologies. Importantly, they will allow for the integration of the diverse datasets necessary to build comprehensive models of the 4D nucleome.

Fourth, genetic, biochemical and biophysical approaches will be developed to measure and perturb the roles of DNA sequences and trans-acting factors (proteins, RNA) in the formation of local and global aspects of the 4D nucleome and their impact on transcription and other nuclear functions.

Fifth, a common vocabulary will be developed to describe nuclear features and biophysically derived principles guiding chromosome folding. This is important because currently different structural descriptions and interpretations have been put forward to describe features detected by different technologies, or even by the same methods. We need better descriptions of the underlying reality of structural features that make up the 4D nucleome, e.g. loops and domains, and develop a consistent terminology as these features are detected by different technologies. This can be achieved by integrated analysis of data obtained with the wide range of technologies employed and under development by the Network. This vocabulary, which will also include nomenclature for non-chromosomal nuclear features, will be of significant value to the community as a whole.

Finally, to enhance synergy between the different groups and prevent the unnecessary duplication of efforts, and to facilitate rapid dissemination of data to the larger scientific community, a shared database and repository will be established which includes all data, detailed protocols, engineered cell lines, and reagents used across the Network. The Network will build and maintain a public browser for all 4D Nucleome data to enable easy access to data and models

## Structure of the 4D Nucleome Network

The 4DN Network encompasses several related efforts (http://www.4dnucleome.org/; Figure 2). First, six Centers make up the Nuclear Organization and Function Interdisciplinary Consortium (NOFIC). Each of these NOFIC Centers will develop genomic and imaging technologies, and implement computational models for understanding the 4D Nucleome. NOFIC centers will also work together with the other components of the Network to benchmark the performance of these experimental and computational tools, and to identify the most appropriate repertoire of methods to study the 4D Nucleome. These studies will be combined with extensive structural and functional validation of observations and models. Ultimately, the NOFIC aims to deliver integrated approaches that can be used towards generating a first draft of a model of the 4D nucleome.

**Figure 2.**
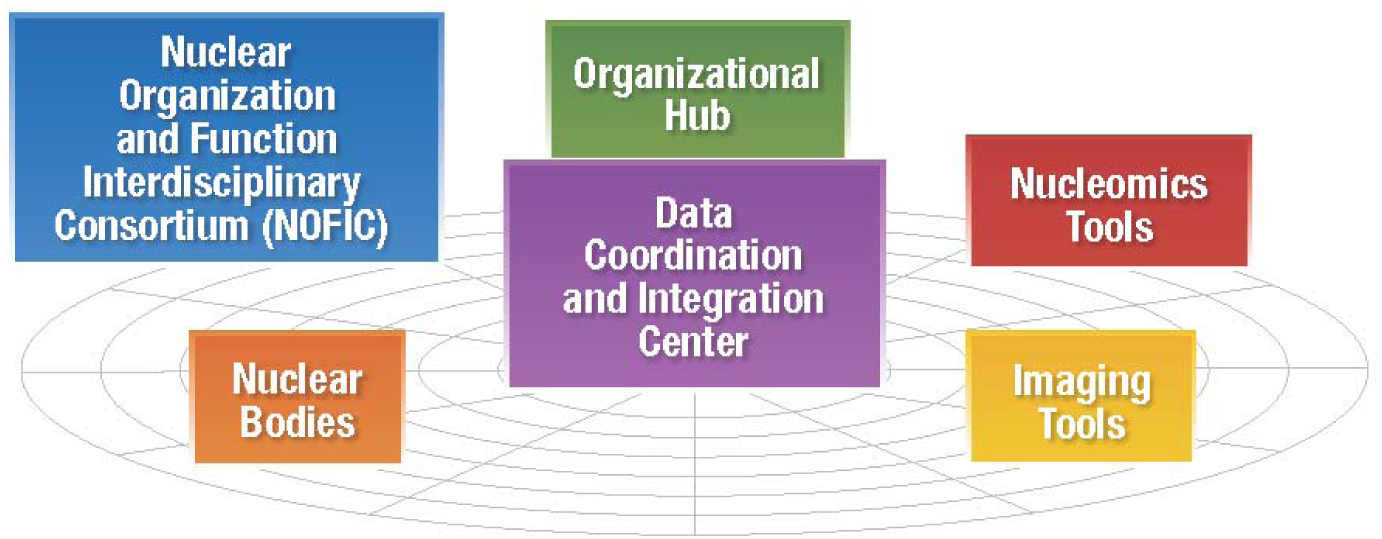
Organization of the 4D Nucleome Network. Six interdisciplinary centers make up the Nuclear Organization and Function Interdisciplinary Consortium. Three additional groups of funded projects focus on analysis of nuclear bodies and development of nucleomics and imaging tools. A Data Coordination and Integration Center collects all data for analysis and public sharing. The Organizational Hub coordinates all Network activities.

Second, ongoing technology development will also be critical to this effort, and is addressed by the 4D Nucleome Network in three ways. (i) New genomic interaction technologies will be developed to study the 4D nucleome at the single cell level, for example, to detect higher-order clusters of interacting genomic loci, to probe the roles of RNA in chromatin architecture, and to engineer new chromatin interactions. (ii) Several groups will develop new imaging and labeling methods to visualize the genome at high resolution, in live cells as well as in tissues, and in relation to genome activity. (iii) New methods will be developed to probe the DNA, RNA and protein composition of subnuclear structures such as the nuclear envelope, the nucleolus and other nuclear bodies. These methods, including procedures to reversibly disrupt these structures, will allow mechanistic studies of the role of nuclear bodies/structures in chromatin organization and gene expression under various conditions of health and disease.

Third, a Data Coordination and Integration Center (DCIC; http://dcic.4dnucleome.org/) will store all data generated by the Network, and coordinate data analysis. The DCIC will maintain a web portal to share data, insights and models with the Network and the larger scientific community. A separate Organizational Hub will coordinate all communication across the Network and organize regular meetings for all participants of the Network.

## Research Plans

### Genomic technologies to reveal the 4D nucleome

Chromosome conformation capture (3C) technologies have been developed to examine long-range interactions across the genome ^15,18^. Genome-wide 3C technologies, *e.g.* Hi-C, have revealed patterns of interaction that define genome structures at various resolutions, including topologically associating domains (TADs) ^16,19,20^. TADs can be hundreds of kilobases in size, often containing several genes and multiple enhancers, at least some of which appear to interact by looping mechanisms. The ChIA-PET method provides finer resolution to detect detailed domain structures defined by chromatin architectural proteins such as CTCF and cohesin, as well as specific enhancer-promoter interactions associated with RNAPII and other TFs ^6^. Furthermore, genome-wide mapping at base pair resolution to detect haplotype specific interactions are in progress, which will enable the connection of chromatin topology to the vast genetic information regarding complex traits and diseases ^17,36,37^. The results generated by these and related methods have been tremendously exciting, and the Network will continue to develop, optimize, and integrate 3C-based technologies with the goal of further elucidating the different levels of chromatin interactions in the nucleus, from the local nucleosome organization of chromatin polymers to their global 3D organization in domains, chromosome territories and in relation to other nuclear structures (Box 1). These efforts will be complemented by orthogonal approaches including direct imaging technologies at both standard and super resolution, X-ray microscopy, and electron microscopy (below and Box 2).

#### Box 1: Genomic technologies currently in use or in development in the 4DN network

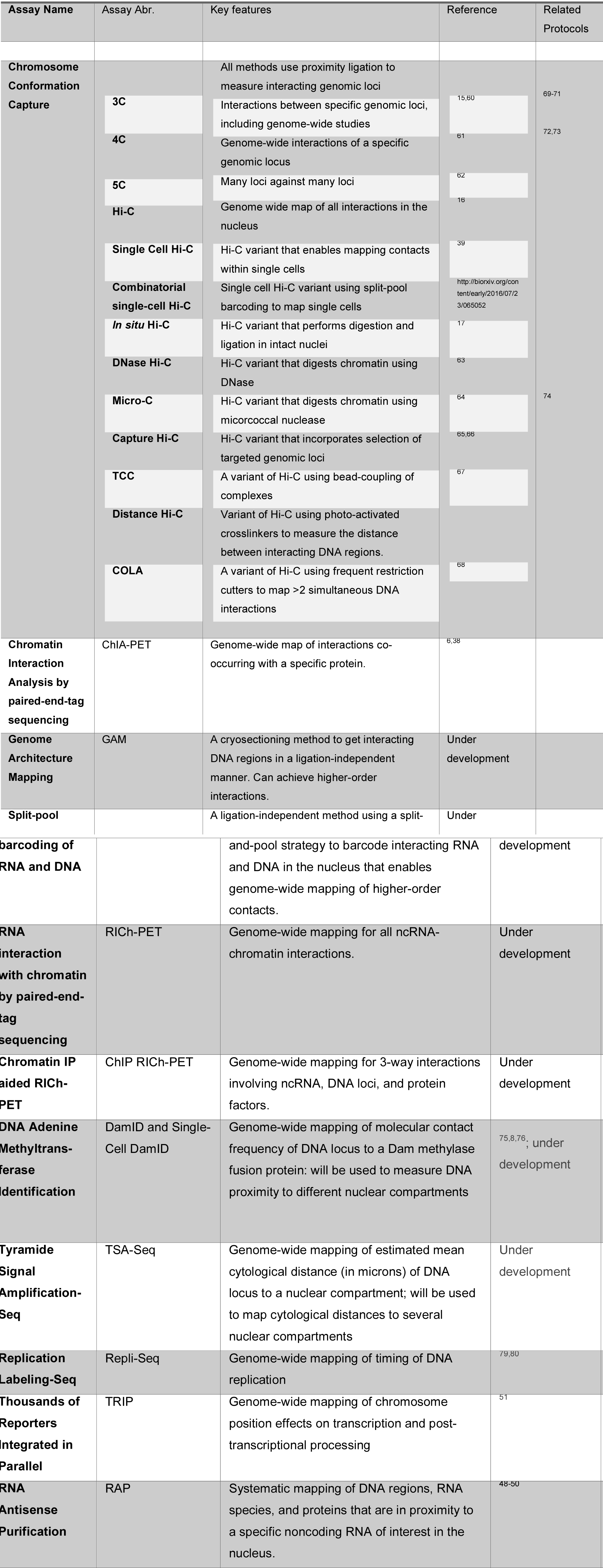

#### Box 2: Imaging technologies currently in use or in development in the 4DN network

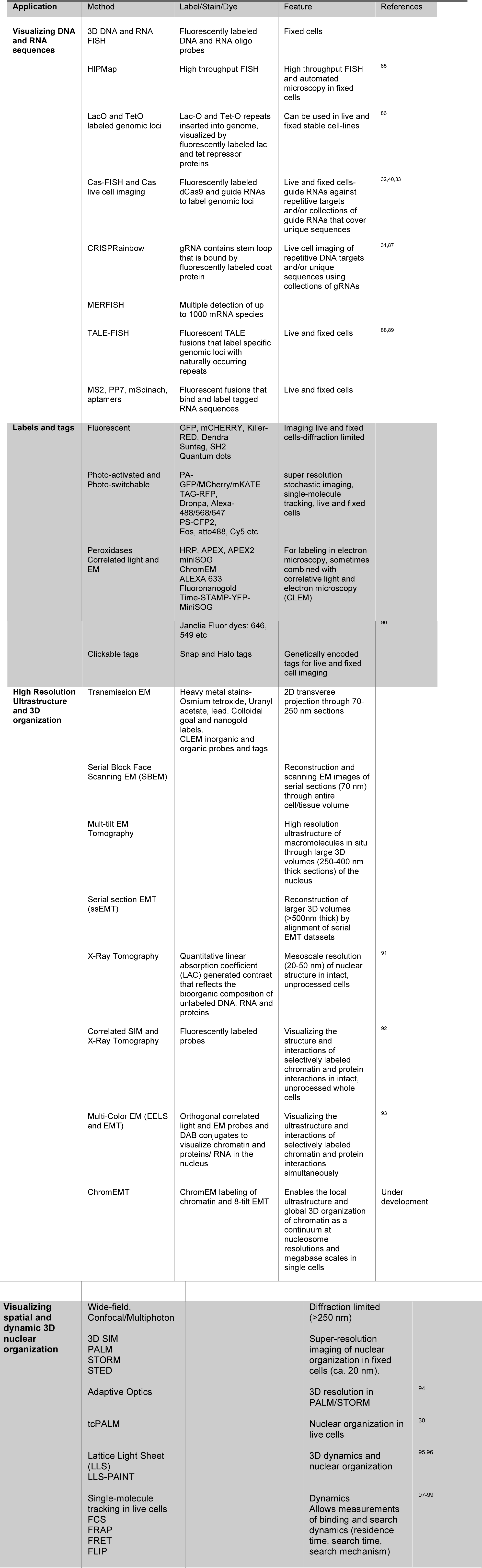

The goal of producing large numbers of data sets of moderate resolution (~1-5 kb) with today’s technology is being complemented by an ongoing effort to invent improved methods for interaction mapping aimed at delivering additional genome interaction information at higher resolution and with haplotype-resolved specificity, lower costs, and at the single cell level. Additional development will also be focused towards implementing genome-wide strategies for mapping RNA-DNA interactions (Box 1).

A major limitation of nearly all 3C methods is that they depend on a single cross-linker, formaldehyde, or on double cross-linking reagents that combine EGS (ethylene glycol bis-succinimidylsuccinate) followed by formaldehyde ^38^. Because of the nature of formaldehyde, which is known to polymerize and crosslink molecules across a large range of distances, this approach complicates the interpretation of the observed chromatin contacts as it lacks precise distance information. The Network will explore novel bivalent photo-activated cross-linkers, which have specificity for DNA and are separated by linkers of defined length and flexibility. Other strategies under development entirely eliminate the use of cross-linkers. Additionally, novel genome-wide methods that enable exploration of higher-order (beyond pairwise) DNA contacts, e.g. genomic DNA regions that form hubs, at various resolutions are being developed. Finally, new methods are being developed that can assess three dimensional genome organization in (thousands of) individual cells ^39^. Clearly, a diversity of existing and new approaches will be necessary to resolve the precise nature of interactions and the physical distances separating the interacting DNAs as well as the cell-to-cell variation in these patterns. The resulting data will provide important information for subsequent modeling of the structural and functional nature of these interactions.

### Imaging the 4D genome

4DN investigators will develop and integrate multi-modal imaging platforms that enable visualizing the dynamics, interactions, and structural organization of the nucleus at unprecedented temporal and spatial resolutions (Box 2). Each of these imaging modalities will have unique and complementary abilities for probing different aspects of genome organization both across the cell cycle and during differentiation. In particular, platforms enabling live cell imaging uniquely allow the dynamics of select chromatin regions and nuclear features to be studied in real time.

Both standard and high-throughput fluorescent in situ hybridization (FISH) using oligonucleotide probes or guide RNA-mediated recruitment of fluorescently labeled dCas9 ("CASFISH" ^33^) in fixed cells will be exploited to image pairwise and multiplexed genomic interactions over different spatial distances in different cell types and states. In addition, CRISPR/dCas9 FISH in live cells ^31,32,40^ and other live cell imaging approaches (Box 2) will be employed to assay the dynamic behavior of particular chromatin regions and/or nuclear structures in real time. These imaging tools will play an important role in benchmarking, validating and complementing data obtained with genomic and proteomic mapping technologies.

New technologies will be developed to exploit chemical, nucleic acid, enzymatic and fluorescent probes to label DNA, RNA, and protein that occur in proximity of specific nuclear bodies. These technologies include HRP-labeled antibodies in TSA-SEQ (Box 1), APEX (ascorbate peroxidase; ^41^) and Killer Red ^42^. Many of these same chemicals and probes are compatible with more established imaging approaches that will also be used to visualize and validate these nuclear body proximity maps. Genome-editing technologies will also be applied to tag a subset of key genomic loci, loops, TADs and potentially newly discovered structures to help visualize these moieties and document their interactions with other nuclear regions in live cells.

Super resolution microscopy, single-molecule tracking techniques and multiplex fluorescent/chemical tags will be applied in living cells to determine the dynamic interactions, diffusion and motion of fluorescently labeled proteins, ncRNAs and genomic loci (Box 2). These live cell imaging approaches are expected to provide novel information regarding search mechanism, binding and residence time of both DNA and protein interactions and will also be relied on extensively to validate and complement genomic methods used by the Network. For example, these methods will be used to determine whether the different contact frequencies of loci in 3D genomic maps reflect their relative distances in real space and/or their interaction frequency and dynamics. Both imaging and genomics groups will focus on particular TADs and loops to define high resolution solutions to the interaction domains of these structures with respect to both other chromatin domains and nuclear bodies using the same cell clones growing under defined conditions.

Soft X-ray Tomography (SXT) will be used to visualize the 3D organization of chromatin in nuclei of cells in the native state (cryo-immobilized). SXT will be used to directly measure chromatin compaction, e.g. in relation to sub-nuclear position, at different stages of cell differentiation and in different cell types. Correlated microscopy approaches will be used to augment ultrastructural data with molecular localization information. Cryogenic Fluorescence Tomography (CFT) will be used to precisely locate molecules in 3D reconstructions of intact, native state cells imaged using Soft X-ray Tomography.

Members of the Network will develop new Electron Microscopy (EM) technologies that enable the local and global structural organization of chromatin to be visualized as a continuum from nucleosome to Mbp scale in both interphase and mitotic cells. One such method, ChromEM, will be combined with new genetic tags and nanoparticle labeling technologies to develop the EM equivalent of ‘multi-color’ fluorescence.

The development of automated imaging analysis pipelines and data standards will be important to extract the maximum structural information possible from these datasets. Further development of software for analyzing, annotating, and archiving imaging data, together with implementation of new approaches for correlating imaging and genomics datasets are major goals of the 4DN Network (see below).

### Nuclear bodies and non-chromatin structures

The nucleus consists of distinct nuclear structures as well as non-chromatin bodies, including the nuclear lamina, nuclear pores, nuclear speckles, and nucleoli ^43,44^. Increasing evidence indicates that specific genomic regions associate with these structures, suggesting that these chromosomal associations may have a functional role in regulating genome function ^8,45,46^.

Major challenges today revolve around our poor understanding and characterization of both the structure and function of these nuclear structures and bodies. Future progress in understanding the relationship between nuclear structure and function demands new methodologies to study the proximity of different chromosomal loci to these different nuclear regions as well as to describe how chromosomal fibers traverse and/or move between these different nuclear regions. Similarly, we envision new technologies to interrogate the protein and RNA content of these different nuclear regions. Additionally, we need high-throughput approaches and imaging methods that query the correlations between genome localization near these different nuclear regions and changes in the functional outputs of these genomic regions. Likewise, we need new methods to dissect the mechanisms by which large genomic regions are targeted to specific nuclear structures / bodies / compartments. Finally approaches to perturb these nuclear structures / bodies /compartments and/or to redirect chromosome regions to different nuclear regions will allow assessing the functional significance of nuclear compartmentalization.

Goals of the 4D Nucleome Network further include development of new mapping methodologies to measure the genome-wide molecular interaction frequency and cytological distance of chromosome loci to major nuclear compartments, including the nuclear lamina, nuclear pores, nuclear speckles, nucleoli, and pericentric heterochromatin (Box 1). Concurrently, new and improved technologies will be developed, including localized APEX (ascorbate peroxidase)-mediated protein biotinylation ^47^, fractionation by cryomilling, and RNA Antisense Purification (RAP) (^48–50^), to catalog and measure the protein and RNA components of these nuclear compartments, as well as both optogenetic and degron-based approaches to alter or disrupt sub-nuclear bodies and compartments. Functional mapping approaches based on Repli-Seq ^9^ and TRIP (thousands of reporters integrated in parallel; ^51^) will provide genome-wide correlations of DNA replication timing and chromosome position effects on transcription and RNA processing that can be correlated with these structural maps (Box 1). New imaging approaches will be developed to correlate chromosome and nuclear compartment dynamics with changes in DNA replication timing, transcriptional activation, and other functional states (Box 2). Computational analyses of these genome mapping data from several cell types will be aimed at identifying possible *cis* and *trans* determinants of nuclear compartmentalization.

### Modeling the 4D Nucleome

As described above, the 4DN network will develop an increasingly powerful repertoire of tools for generating high-throughput data that inform us about 3D/4D nuclear architecture. The major classes of methods yield data that are informative with respect to biochemically-derived contact probabilities of loci (e.g. 3C and its derivatives) or the spatial distances between loci (e.g. FISH and other imaging methods). Additional technologies yield data that are informative of other aspects of nuclear architecture (e.g. the relative positions and contents of nuclear structures in relation to one another as well as the folded genome).

In parallel and often in concert with the emergence of these experimental methods has been the development of computational approaches for modeling the spatial organization of the genome. There are at least two major computational approaches for global modeling of genome architecture on the basis of experimental data: *data-driven* and *de novo* approaches ^34,35^. *Data-driven approaches* directly use experimental data to produce a single conformation or an ensemble of conformations that best match an experimentally observed set of contact probabilities or distances ^52^. *De novo* modeling, on the other hand, produces ensembles of conformations that result from known or hypothesized physical or biological processes, and tests whether these ensembles are consistent with features of experimental contact frequency maps and imaging data. Such *de novo* models can suggest specific molecular mechanisms and principles of chromosome organization, thus going far beyond the experimental data ^35^.

Although these approaches to modeling genome architecture have been useful for making sense of the observed data and relating it to principles of chromosome organization, it is clear that we are still far from a high-confidence, high-resolution map of genome architecture. Therefore, an overarching goal of the Network is to generate reliable and validated structural maps representing the 4D Nucleome for different biological states of the genome.

A number of challenges and opportunities with respect to computational modeling of genome architecture will be addressed. First, computational models of global genome architecture have typically relied on a single data type, e.g. Hi-C, which limits the power of such analyses because it is likely that Hi-C or ChIA-PET or any other one method does not capture all aspects of the 4D nucleome and does not sufficiently constrain models. The Network will apply a wide diversity of technologies, and each of these is likely to capture complementary aspects of genome organization, e.g. contacts by Hi-C and distances by imaging ^53,54^. A major challenge is thus to adapt current modeling approaches so that data obtained with these methods (e.g. Hi-C and imaging data) can be systematically integrated and used to generate comprehensive structural and dynamic models of the 4D nucleome. Second, nearly all current genomic methods yield data that are derived from an ensemble of thousands to millions of cells, obscuring structural heterogeneity that exists among single cells. A number of groups within the Network are developing methods for generating global data from large numbers of single cells, which will present new computational challenges for integrating with current modeling approaches ^39^. It is possible that some of these methods will yield functional data on these same single cells (i.e. Hi-C and RNA-seq, from each of many single cells), which would represent a significant opportunity for directly relating structure to function. Third, most current models ignore the fact that mammalian cells are diploid, i.e. they do not distinguish or separately model homologous chromosomes, which will be particularly important for modeling based on single-cell data. It is critical to develop experimental methods and computational pipelines that overcome this deficiency ^17,36,37^; the haplotype-resolution of the genomes of the common cell types chosen by the consortium will aid in this goal. Fourth, contemporary approaches for modeling genome architecture are typically static rather than dynamic, reflecting the static nature of available Hi-C data and the majority of imaging data. As we are increasingly able to visualize (via direct imaging) or infer (via single cell or bulk Hi-C analyses of time series, live cell imaging) dynamic processes, e.g. differentiation, response to physiological stimuli, and cell cycle progression, it will be essential that these observations can be integrated into computational models. Fifth, new data and future models should help to connect genome architecture and other aspects of genome function, e.g. by suggesting molecular mechanisms of how transcription factor binding or epigenetic modifications can lead to formation of active/inactive chromatin compartments ^55,56^, etc. The inferred relationships between these features and genome architecture would serve as a means of generating testable predictions of sequence-structure-function relationships, i.e. how nuclear architecture relates to nuclear function.

### Perturbation and manipulation of the 4D Nucleome to relate structure to function

A critical and overarching goal is to determine how genome structure and chromatin interactions at multiple scales modulate genome function in health and disease. However, despite significant progress, many of the mechanisms by which chromosome structures such as chromatin domains are formed and how they are related to local genome activity remain poorly understood.

Therefore, the 4DN Network will explore a range of experimental approaches to specifically manipulate and perturb different features of the 4DN. First, using CRISPR/Cas9 technologies and targeted DNA integration experiments, DNA elements involved in specific chromatin structures, e.g. domain boundaries or chromatin loops, can be altered, re-located or deleted ^57,58^. Second, defined chromatin structures such as chromatin loops will be engineered de novo by targeting proteins that can (inducibly) dimerize with their partner looping proteins (e.g. ^7^). Third, other CRISPR/Cas9 approaches will be used to target enzymes (e.g. histone modifying enzymes, structural proteins) or ncRNAs to specific sites in the genome. Fourth, several groups will perturb nuclear compartmentalization by developing methods for “rewiring” chromosome regions to different nuclear compartments, either by integrating specific DNA sequences that are capable of autonomous targeting the locus to different nuclear compartments or bodies or by tethering certain proteins to these chromosomal loci to accomplish similar re-positioning. Fifth, cell lines will be generated for conditional/temporal ablation of nuclear bodies or candidate chromosome architectural proteins (such as CTCF and cohesin) or RNAs. Sixth, additional methods will be developed to nucleate nuclear bodies at specific chromosomal loci. Finally, biophysical approaches will be developed to micro-mechanically perturb cell nuclei and chromosomes followed by direct imaging of specific loci ^59^. Analysis of the effects of any of these perturbations on processes such as gene expression and DNA replication will then provide deeper mechanistic insights into the roles of chromosome structure and nuclear organization in regulating the genome.

### Data sharing and standards

An important goal for the Network will be to develop practical guidelines for data formats, metadata (detailed descriptions of how the data was acquired), standards, quality control measures, and other key data-related issues. Another goal will then be to make this production-quality data rapidly and readily accessible, both within the Network and the entire scientific community. These efforts will be of particular importance for new technologies for which standards for sharing data and assessing data quality have not yet been established. Such standards will greatly enhance the usefulness of the datasets for the broader scientific community beyond those who generate the data.

For sequencing-based technologies, data format standards to represent sequences and alignments have long been present (e.g. fastq, bam/sam). However, common formats to represent three-dimensional interactions are yet to be developed. These formats need to account for large data sizes and the constraints imposed by different computer architectures. For Hi-C data, for example, the genome-wide contact probability map is an N^2 matrix, where N is the spatial resolution (for 10kb resolution, N=300,000), with most of the entries being empty; whereas for ChIA-PET data the detected chromatin contacts are between selected loci bound by a protein of interest, which greatly reduces the complexity of the interaction matrix. There are multiple ways to represent such sparse matrices, appropriate for different analysis and storage approaches. For imaging technologies, the situation is even more challenging, as the types of microscopes employed are highly variable, and the data formats and analysis tools are often manufacturer-dependent. Standards to unify data and metadata from different manufacturers, such as the Open Microscopy Environment (https://www.openmicroscopy.org/site), are under development. These standards also need to accommodate the rapid and ongoing developments in super-resolution microscopy technologies.

A related issue is to define a set of appropriate metadata fields and minimum metadata requirements such that sufficient and useful details are available to other investigators outside the Network. While it is acknowledged that not all information can be captured about an experiment, collecting pertinent information (without unnecessarily burdening the experimentalists) will increase the reproducibility of experiments and the likelihood that the data will be re-used by other investigators. Towards this end, the 4DN Consortium has established formal working groups, including the 4DN Data Analysis Group, Omics Standards Group and Imaging Data Standards Group, which will play critical roles in defining these 4DN standards and data analysis protocols.

Developing a set of measures for assessing data quality and determining appropriate thresholds will be important for ensuring high-quality 4DN data. An important measure of data quality and reliability of new technologies is the reproducibility of results between repeated experiments. Reproducibility can be assessed at multiple levels: broadly speaking, technical reproducibility measures how well a technique performs for the same starting material, whereas biological reproducibility should also capture all other variations including heterogeneity among samples. Data quality standards and thresholds for what constitutes ‘good’ data will change over time as technologies improve and analysis methods are refined. To accommodate this fluidity, the 4DN Analysis Group will compute and make available relevant quality control measures and provide recommendations on expected quality standard thresholds so that investigators can make decisions regarding the utility of specific datasets for addressing their specific questions.

### Outlook

After determining the complete DNA sequence of the human genome, the variations in this sequence in individuals and across populations, and mapping most genes and potential regulatory elements, we are now in a position that can be considered the third phase of the human genome project. In this phase, the spatial organization of the genome is elucidated and its functional implications revealed. This requires a wide array of technologies from the fields of imaging, genomics, genetic engineering, biophysics, computational biology and mathematical modeling. The 4D Nucleome Network, as presented here, provides a mechanism to address this uniquely interdisciplinary challenge. Further, the policy of openness and transparency both within the Network and with the broader scientific community, and the public sharing of all methods, data and models will ensure rapid dissemination of new knowledge, further enhancing the potential impact of the work. This will also require fostering collaborations and establishing connections to other related efforts around the world that are currently under development.

Together these highly integrated studies, both within the 4DN Network, and with partners around the world, promise to allow moving from a one-dimensional representation of the genome as a long DNA sequence to a spatially and dynamically organized three dimensional structure of the living genome inside cells. This structure will place genes and regulatory elements that can be far apart along the DNA, in a three-dimensional and dynamic context, revealing their functional relationships and providing a new understanding of how the genome is regulated.

